# FREQ-NESS Reveals the Dynamic Reconfiguration of Frequency-Resolved Brain Networks During Auditory Stimulation

**DOI:** 10.1101/2024.08.28.610155

**Authors:** M. Rosso, G. Fernández-Rubio, P. Keller, E. Brattico, P. Vuust, M. L. Kringelbach, L. Bonetti

## Abstract

The brain is a dynamic system whose network organization is often studied by focusing on specific frequency bands or anatomical regions, leading to fragmented insights, or by employing complex and elaborate methods that hinder straightforward interpretations. To address this issue, a new analytical pipeline named *FREQuency-resolved Network Estimation via Source Separation* (FREQ-NESS) is introduced. This is designed to estimate the activation and spatial configuration of simultaneous brain networks across frequencies by analyzing the frequency-resolved multivariate covariance between whole-brain voxel time series. FREQ-NESS is applied to source-reconstructed magnetoencephalography (MEG) data during resting state and isochronous auditory stimulation. Results reveal simultaneous, frequency-specific brain networks during resting state, such as the default mode, alpha-band, and motor-beta networks. During auditory stimulation, FREQ-NESS detects: (1) emergence of networks attuned to the stimulation frequency, (2) spatial reorganization of existing networks, such as alpha-band networks shifting from occipital to sensorimotor areas, (3) stability of networks unaffected by auditory stimuli. Furthermore, auditory stimulation significantly enhances cross-frequency coupling, with the phase of attuned auditory networks modulating the gamma band amplitude of medial temporal lobe networks. In conclusion, FREQ-NESS effectively maps the brain’s spatiotemporal dynamics, providing a comprehensive view of brain function by revealing simultaneous, frequency-resolved networks and their interaction.

## Introduction

The ultimate goal of cognitive neuroscience is to understand how the human brain functions as an interface with the external world ^1^. This endeavor involves characterizing the brain’s intrinsic dynamics ^2–4^ and their interaction with continuous streams of information sampled from the environment ^5–8^. Despite significant progress, this pursuit presents two critical challenges that are not fully resolved yet.

The first challenge lies in understanding the functional organization of the brain. There is wide consensus that optimal functioning is supported by the coordination of functionally specialized networks ^9^. Accurately capturing the spatiotemporal organization of these networks, their intrinsic dynamics, and their complex interactions remains a cornerstone of cognitive neuroscience. Despite substantial progress, debates persist regarding the most effective methods for analyzing and interpreting brain networks in different experimental scenarios. The second challenge involves disentangling the multitude of concurrent neural processes underpinned by such networks, which overlap in time and space regardless of whether the brain is at rest or engaging with external stimuli ^10,11^. This overlap generates mixed neurophysiological signals, hindering the isolation of the underlying neural mechanisms and their contribution to cognition and behavior.

Addressing these challenges requires defining a criterion to separate overlapping neurophysiological signals, ideally through multivariate solutions that leverage the multiple recording units characteristic of neurophysiological data ^11^. A key criterion for distinguishing such overlapping signals is neural frequency, both at rest and during tasks ^12–17^. As a matter of fact, strong associations have been reported between oscillatory brain activity and cognitive processing ^4,14,18,19^. Similarly, atlas-based approaches suggest that brain regions exhibit dominant frequency modes ^20,21^, reinforcing the notion of a functional link between temporal and spatial organization. Based on this evidence, Jiang et al. ^22^ recently introduced a multivariate approach to analyze large-scale spatial–rhythmic networks (SRN) with distinct spatiotemporal dynamics. Applying this method to source-reconstructed EEG data projected onto anatomical ROIs, the authors provided valuable insights into cognitive functions and disease by implementing an adaptive frequency decomposition of the signal.

Taken together, these findings suggest that frequency is a key criterion for disentangling overlapping signals and identifying functionally distinct brain networks. However, an essential question remains: how should neural processes be effectively separated on an analytical level? As mentioned, multivariate approaches offer a powerful means to isolate concurrent brain processes that overlap in time and space, as supported by extensive research ^23–27^. Yet, current methods face critical limitations.

Many multivariate approaches rely on predefined, anatomical parcellations in regions of interest (ROIs) to simplify computations and reduce complexity. In particular, ROI-based methods inherently follow a hypothesis-driven framework, requiring researchers to select regions in advance and therefore underutilize the richness of neural recordings. We argue that a voxel-based approach better leverages the full spatial complexity of the data, offering three key advantages. First, it preserves spatial gradients of activation within and across parcels. Second, it mitigates parcellation biases by avoiding artifacts caused by activity at parcel boundaries and inconsistencies resulting from variable parcel dimensions. Third, it minimizes assumptions by enabling fully data-driven analyses that preserve the intrinsic structure of neural recordings and enhance the characterization of frequency-specific networks.

Another widely used approximation concerns the frequency domain itself. In particular, studies examining functional connectivity often focused on canonical and relatively broad frequency bands (e.g., delta, theta, alpha, beta, gamma) ^27–29^. Although informative, these bands lack the granularity necessary for fully frequency-resolved analyses. Greater resolution can be achieved by targeting specific frequencies of interest or by scanning a dense sample of frequencies across the spectrum, to uncover subtle yet meaningful differences ^30^. Moreover, although it is common practice to estimate brain networks solely based on filtered data, there are analytical tools which are specifically designed to isolate the contribution of frequency- specific components against the multivariate broadband signal ^11,24^. Thus, these tools are conceptually and mathematically well-suited to our goals, providing an optimal opportunity for examining the frequency-specific properties of brain networks.

A prominent example is generalized eigendecomposition (GED) ^11^. Due to its flexibility in defining an optimization criterion for source separation, GED offers a superior solution for studying frequency-related signal content compared to several other methods including different linear decomposition techniques such as principal component analysis (PCA) or independent component analysis (ICA)^11,28,29^. Among other applications ^11,31,32^, GED can decompose multivariate signals into components exhibiting activity at a defined frequency that characterizes the process of interest ^26,32–36^. This feature makes it an ideal framework to investigate the brain in resting state and its processing of naturalistic stimuli such as music and speech as they continuously unfold over time. This occurs because the temporal structure of these stimuli elicits neural activation at matching frequencies ^37–39^, and the associated domain-specific brain networks are characterized by unique spectral fingerprints ^26,33,37–40^. While previous applications of GED have demonstrated its reliability in separating sources related to distinct cognitive processes within specific frequency bands, its contributions to understanding the spatial extent of these processes and the interactions among the underlying brain networks operating at different frequencies have been limited. This limitation primarily arose from the predominant application of GED to sensor data matrices in non-invasive neurophysiology (e.g., ^26,33–36,40–43)^. Although valuable, this approach provided limited spatial information about the distinct contributions of different brain regions and networks to the various cognitive processes inferred from multivariate neurophysiological data.

Building on prior work, the current study aims to characterize the spatiotemporal organization of brain networks and disentangle overlapping neural processes through frequency specificity, directly addressing the challenges of brain organization and concurrent neural processes discussed earlier. Specifically, we applied a refined analytical pipeline, termed *FREQuency-resolved Network Estimation via Source Separation* (FREQ-NESS), which uses GED on source-reconstructed MEG data in 8mm³ space in a frequency-resolved manner. By integrating well-established analytical tools, FREQ-NESS enhances traditional approaches in three critical ways. First, it optimizes the separation of brain networks based on their frequency content, quantifying the prominence of each frequency-specific network over the background broadband activity. Second, it produces spatial activation patterns with fine granularity, preserving the full spatial complexity of the data while avoiding the constraints related to ROI-based methods. Third, it generates activation time series for each frequency-specific brain network, facilitating further analysis of within-network dynamics as well as between-network interactions via cross-frequency coupling. As a result, it captures the global organization of the brain in the form of a frequency-specific network landscape, characterizing their spatial and temporal extents at rest and its reconfiguration during cognitive processes.

In this study, FREQ-NESS enabled us to test specific hypotheses on the functional organization of the brain within a minimalistic experimental paradigm involving resting state and rhythmic auditory stimulation. Initially, we applied this methodology during resting state aiming to concurrently capture its underlying brain networks (e.g., default-mode network, alpha-band network, beta-band network, gamma-band network ^20,44^). Subsequently, we recorded participants’ brain activity during rhythmic auditory stimulation at a defined frequency to investigate how their brain networks were reshaped by such low-effort task.

First, we hypothesized that diverse functional brain networks would be simultaneously active during resting state, each characterized by distinct frequency and spatial profiles, with varying degrees of prominence relative to one another. Second, we expected that the overall composition of the resting-state network landscape would be significantly reshaped in response to isochronous auditory stimulation, altering the relative prominence and spatial distribution of specific networks. Third, we hypothesized that, during basic auditory perception, coupling would emerge between the stimulation frequency and higher frequencies, linking the auditory network with a distributed network of brain regions.

In conclusion, FREQ-NESS addresses previous literature gaps by providing a refined method for studying functional brain networks and their interactions. By applying this pipeline to both resting state and auditory perception data, we reveal new insights into the dynamic organization of the human brain as a landscape of frequency-specific brain networks and its reconfiguration in response to environmental interactions. Furthermore, we address its effectiveness and versatility for adoption across a wide range of experimental scenarios in human neuroscience.

## Results

### Experimental design

Twenty-nine (n = 29) participants underwent five minutes of MEG recording in two conditions: resting state (RS) and passive listening (PL) to an isochronous auditory tone sequence stimulus presented at a rate of 2.4 Hz (**Figure 1a** and **b**). Two participants did not undergo the PL recording, and one participant was discarded due to technical issues during the data collection, leaving us with twenty-six (n = 26) participants eligible for within-subject comparisons in both conditions.

**Figure 1:**
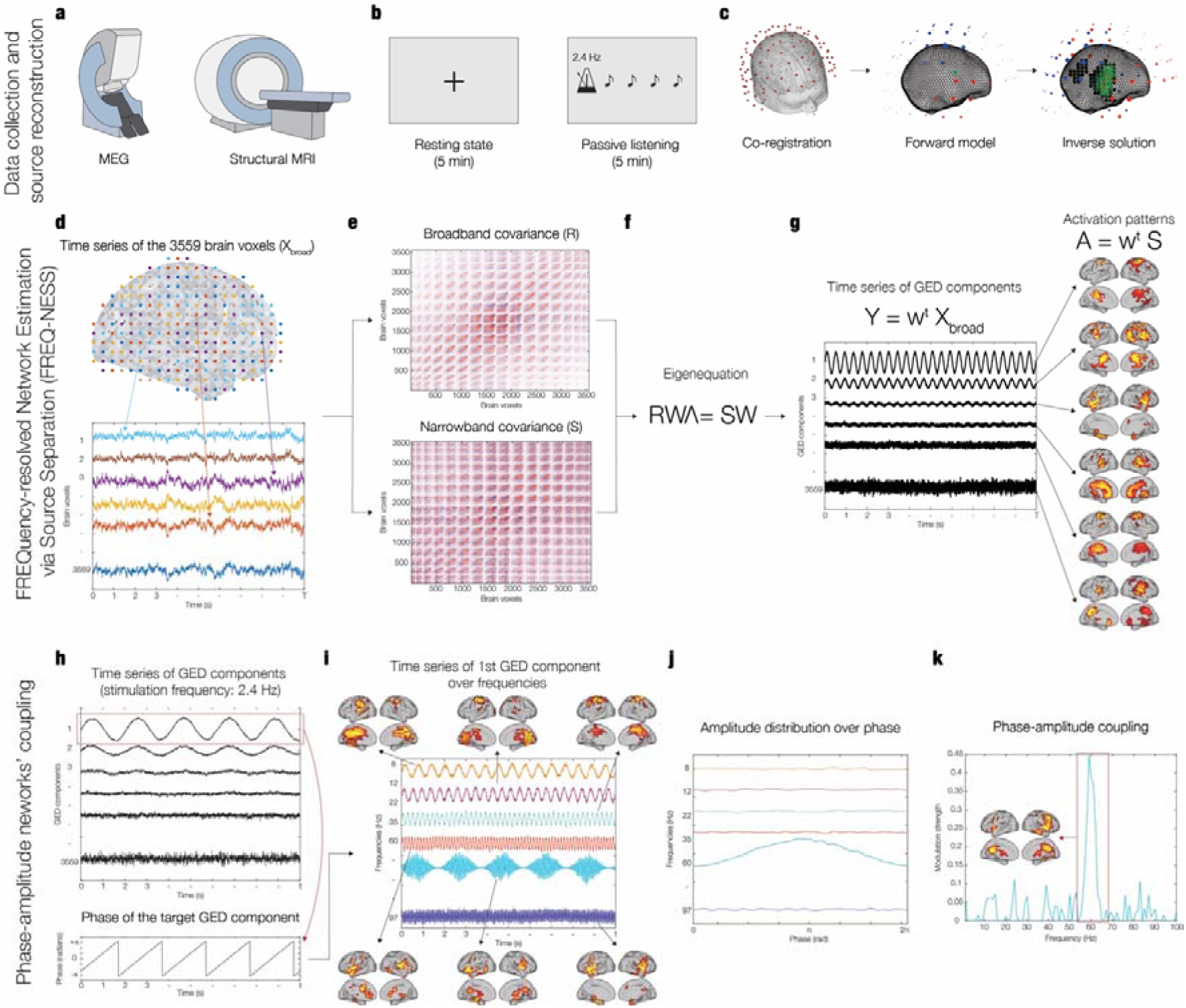
FREQuency-resolved network estimation via source separation (FREQ-NESS). The figure provides an overview of the FREQ-NESS analytical pipeline. **a -** MEG was used to collect neurophysiological data in the experimental conditions, while structural MRI was acquired in a separate session. **b -** Participants underwent MEG recordings during two experimental conditions: resting state (5 minutes) and passive listening to isochronous auditory tone sequence (5 minutes) at a rate of 2.4 Hz. **c –** To compute MEG source reconstruction, we performed co-registration between MEG data and individual MRI anatomical scans, forward model (single shell), and inverse solution using beamforming. **d –** The source reconstruction generated one time series for each of the 3559 brain voxels from an 8-mm grid parcellation of the brain. **e -** The covariance matrices R and S were computed from the broadband voxel data matrix (R) and from the narrowband filtered voxel data matrix (S); the procedure was repeated for a sample of 86 frequencies, computing a new S covariance matrix upon each iteration. **f -** Generalized eigendecomposition (GED) was computed by solving the eigenequation RWΛ = SW for the eigenvectors (W) to find the weighted combinations of brain voxels to separate frequency-specific networks; the associated eigenvalues (Λ) express the amount of variance explained by each network component. **g -** Network activation time series and spatial patterns were computed. The spatial filters (W) were applied to the broadband voxel data matrix (X**_broad_**) via matrix multiplication to reconstruct its components (Y) as network activation time series; the same spatial filters were applied to the narrowband covariance matrix S to compute the spatial activation patterns (A) as the components’ projections in voxel space. **h -** The phase of the bandpass filtered time series of the 2.4 Hz components of interest (brain networks) is extracted via Hilbert transform for subsequent cross-frequency coupling (CFC) analysis. **i -** To investigate the modulation of network interactions (phase of the 2.4 Hz brain networks modulating the amplitude over all the frequency bins), the modulation of frequency-specific brain networks as a function of the low-frequency modulating network was quantified as amplitude modulation. **j -** For each network, mean power values were calculated for all the phase bins, resulting in an amplitude distribution over the 2.4 Hz network’s phase. **k –** For each frequency-specific network, an index of modulation strength was computed by fitting a sinewave to the modulation curves and computing its amplitude; this enabled the identification of significant interactions across networks operating at specific frequencies (here, phase of the 2.4 Hz brain networks modulating the amplitude over all the frequency bins).

FREQ-NESS was used to characterize the temporal and spatial configuration of brain networks operating at specific frequencies, and to investigate their rearrangement as a result of continuous auditory stimulation. To this end, we performed GED on the entire brain voxel data matrix as reconstructed via beamforming. We iterated the procedure for both conditions, for each participant, and for the entire frequency spectrum from 0.2 to 97.6 Hz, in intervals of 1.2 Hz (for a similar scanning procedure, see also ^30^). For the sake of consistency with existing literature, we will often refer to the canonical division in frequency bands when reporting our results ^45^: delta (0.2 – 4 Hz), theta (4 – 8 Hz), alpha (8 – 13 Hz), beta (13 – 30 Hz), and gamma (30 – 97.6 Hz). However, our primary focus is on specific frequencies within these ranges.

For each frequency, we computed the eigenvectors to decompose the multivariate signal in its components carrying the most prominent narrowband activity at the specified frequencies, as contrasted with broadband activity. The same eigenvectors were used to filter the covariance matrix of narrowband activity, computing the spatial activation patterns of the respective component. Given the dataset consisted of 3559 voxels time series, 3559 components were extracted and sorted in descending order by explained variance. This resulted in a network landscape consisting of the eigenvalue distribution over the frequency spectrum and the associated spatial activation patterns. The associated network activation time series were used at a subsequent stage to perform cross-frequency coupling analysis between networks.

### Variance explained by the brain networks in resting state and passive listening

To assess changes in the network landscape associated with the auditory stimulation in PL against RS, we conducted a series of Wilcoxon signed-rank tests. Considering the exponential decay observed for the eigenvalues returned by GED, we focused our statistics on the top 10 eigenvalues (i.e. the eigenvalues corresponding to the 10 GED components which explained the highest variance). We contrasted PL vs RS over the entire frequency spectrum and applied False Discovery Rate (FDR) correction for multiple comparisons. In this context, every normalized eigenvalue expresses the percentage of total variance explained by the respective signal components in GED space ^44,45^. Positive z-values emerging from the Wilcoxon test were interpreted as the emergence of frequency-resolved networks of PL against the RS baseline, whereas negative z-values were interpreted as their suppression. To increase readability of the results, we report statistical tests for only the top three eigenvalues up to 40.9 Hz in **Figure 2**, while detailed depiction of full statistical test outcomes for the full range (top 10 eigenvalues over the spectrum up to 97.6 Hz) is provided in **Figure S1**.

**Figure 2:**
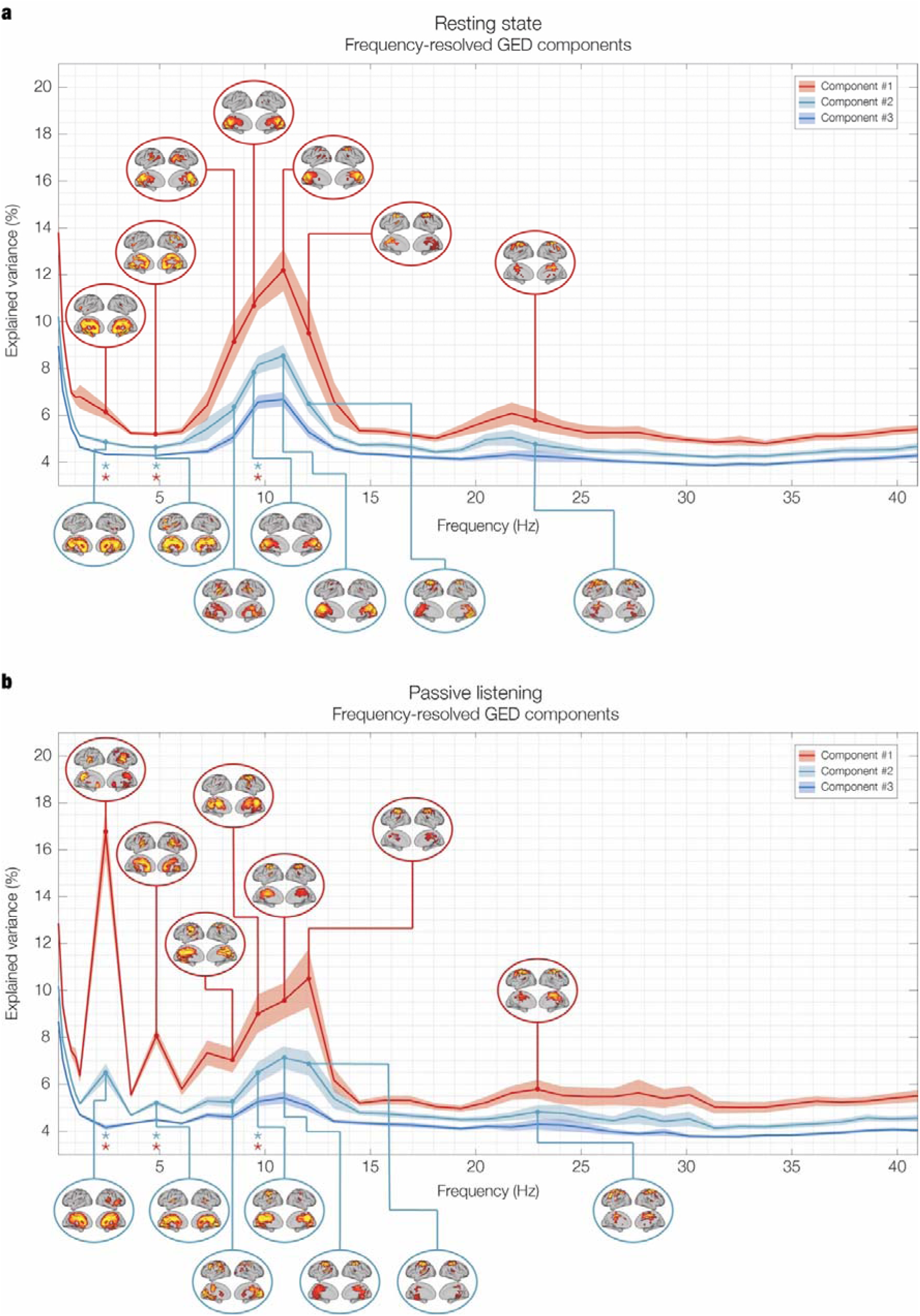
Network landscape: eigenspectrum and spatial activation patterns during rest and passive listening. This figure illustrates the frequency-resolved brain networks estimated via FREQ-NESS in two experimental conditions: (a) resting state and (b) passive listening. The first element of the network landscape is the **eigenspectrum**, which consists of the eigenvalues expressing the percentage of variance explained by the associated GED components, plotted over the frequencies. The solid lines represent the mean normalized eigenvalues averaged across 26 participants (N = 26), with shaded areas indicating the standard error of the mean (SEM). Stars denote frequency bins where significant differences were detected using a two-tailed signed-rank Wilcoxon test, following FDR correction for multiple comparisons. The second element displays the **spatial activation patterns** associated with the top GED components for the frequencies exhibiting the highest explained variance. Each pattern highlights the extent of each brain voxel’s contribution to the respective network. To improve visualization, the eigenspectra are shown here for the first three GED components (sorted by explained variance) over a portion of the frequency spectrum (0.2 to 40.9 Hz), while the topographies of the first two components are shown for the frequencies with the highest explained variance. For an extended version of the network landscape, see **Figure S1**. **a - Resting state (RS)**: The network landscape reveals a 1/f exponential decay of explained variance in the low delta range and local maxima in the alpha (10.9 Hz) and beta (22.9 Hz) frequency ranges. The spatial activation patterns show the typical topographies expected from the brain at rest, namely the broad mesial distribution associated with the DMN for low frequencies not engaged in stimulus processing (2.4 Hz and 4.8 Hz), parieto-occipital distribution for the alpha network (10.9 Hz), and sensorimotor involvement for the beta network (22.9 Hz). Notably, while frequencies on the left side of the alpha peak exhibit parieto-occipital activation (8.4 Hz and 9.6 Hz), the higher end of the range is characterized by sensorimotor activation (12.1 Hz). In the beta range, the explained variance peaks at 22.9 Hz, and the spatial pattern shows activation limited to sensorimotor regions. **b – Passive Listening (PL)**: Significant reorganization of brain networks occurs in response to auditory stimulation. The eigenspectrum exhibits a pronounced peak at the stimulation frequency of 2.4 Hz and its first harmonic at 4.8 Hz, reflecting the emergence of brain networks attuned to the rhythmic auditory stimulus. The spatial activation patterns at 2.4 Hz show focal engagement of the right auditory cortex, particularly in Heschl’s gyrus, with additional involvement of broader medial temporal regions in the 4.8 Hz component, indicative of secondary auditory processing. Another key change observed in this condition is the shift in the alpha peak from 10.9 Hz in the resting state to 12.1 Hz during listening, accompanied by a posterior-to-anterior shift in spatial activation from parieto-occipital to sensorimotor regions. This reorganization suggests that auditory stimulation induces a rearrangement of alpha networks, likely reflecting increased readiness for action and sensory processing in response to the rhythmic stimulus. The arrangement of beta remains stable across conditions, both in terms of eigenspectrum and spatial patterns.

Significant changes in the first two eigenvalues were found for the following frequencies: 2.4 Hz (1^st^ component: *p* < 0.001, FDR-corrected; 2^nd^ component: *p* < 0.001, FDR-corrected), 4.8 Hz (1^st^ component: *p* < 0.001, FDR-corrected; 2^nd^ component: *p* < 0.001, FDR-corrected), and 9.6 Hz (1^st^ component: *p* < 0.001, FDR-corrected; 2^nd^ component: *p* < 0.001, FDR-corrected).

The prominent peaks at 2.4 Hz indicate a strong attunement of the brain to the frequency of the auditory stimulation. By ‘attunement’, we mean that the spectral content of a network component becomes highly concentrated around a specific frequency as a direct result of the stimulation. The term is conceptually and operationally different from ‘entrainment’ as the latter assumes a specific phase-based mechanism leading to the match in the stimulation frequency ^46,47^. Whilst a possibility ^33,48^, we do not claim that entrainment is the mechanism underlying our result. As a qualitative observation, we note that this attunement also reshaped the 1/f exponential decay on the left end of the spectrum. Specifically, we observed a suppression of the aperiodic components of the brain signal in the background ^49^, caused by the emergence of primary and secondary frequency-resolved auditory networks. Based on the harmonic relationship with the stimulation frequency and the associated spatial activation pattern, we interpret the emergence of the 4.8 Hz as attunement of auditory networks alongside the 2.4 Hz components.

The significant change at 9.6 Hz is the result of a rearrangement in the alpha region of the landscape. Specifically, the peak of the top components shifted from 10.9 Hz in RS to 12.1 Hz in PL, biasing the symmetry of the Gaussian-shaped distributions observed for RS to the right. The result indicates an imbalance between alpha ranges across conditions, with auditory stimulation resulting in a suppression of the lower and an enhancement of the higher alpha range. We interpret this finding as indicative of a re-attunement of the alpha networks during rhythmic auditory processing.

Beta and gamma ranges did not exhibit any significant change between experimental conditions in terms of explained variance. Detailed statistical results for all components over the frequency spectrum are reported in **Table S1**.

In order to complement the original FDR-corrected statistical tests, we conducted an additional analysis using a cluster-based permutation test to identify significant clusters of consecutive frequency bins across the spectrum. This analysis revealed a significant low-alpha cluster spanning 7.2 Hz to 10.9 Hz for the first two GED components (cluster size = 4 consecutive frequency bins, p-value < 0.001), providing further support for our conclusion that alpha networks shift their activity toward higher frequencies during auditory stimulation. However, as expected, no significant clusters were detected at the stimulation frequency (2.4 Hz) or its first harmonic (4.8 Hz), even though these frequencies showed the largest differences from the resting state baseline and the smallest p-values (see **Figure 2** and **Table S1**). This outcome highlights an inherent limitation of cluster-based permutation tests, which require consecutive bins to form clusters. Since the 2.4 Hz and 4.8 Hz networks exhibit high frequency specificity with sharp drops in explained variance in neighboring bins, they do not meet the clustering criterion. While cluster-based permutation tests are effective for managing multiple comparisons and identifying broader, contiguous frequency clusters, these results highlight their reduced sensitivity to localized, frequency-specific effects.

After detecting statistically significant changes in the variance explained by the GED components across experimental conditions, we computed the spatial activation patterns to provide complementary qualitative information, as typically done in multivariate methods for neuroimaging (see ^11,23,33^). As shown in **Figure 2** and described in detail in the Methods section, spatial activation patterns were computed for each eigenvalue associated to the GED components (i.e. brain networks) to produce their topographical projection in brain voxel space. Please note that in all the figures in this article, only the activation patterns associated to the first two components of the most relevant frequencies are shown. For each activation pattern, only the voxels whose activation exceeded their mean by more than one standard deviation were displayed on the brain template (see ^50^).

The entire delta range (0.2 – 4 Hz) in RS was characterized by the spatial distribution typical of the default-mode network (DMN) ^44^. At 2.4 Hz, in absence of stimulation, both the first and the second component matched this configuration. Conversely, in PL, the first component involved primarily the right auditory cortex with a major focus on the Heschl’s gyrus ^51^, while the second component recruited a larger brain network previously connected to secondary auditory processing ^50,52^ and including regions of the medial temporal lobe such as hippocampal regions, inferior temporal cortex, insula, ventromedial prefrontal cortex, as well as the cingulate gyrus. Taken together, these spatial changes show the unequivocal involvement of early and late stages of auditory processing, marking a transition from DMN to auditory networks associated to the frequency attunement. This physiological interpretation is corroborated by the observation that, over the whole delta range, both the frequency attunement and the spatial clustering during PL occurred prominently and selectively for the stimulation frequency.

In the theta range (4-8 Hz), a similar pattern was observed for the first harmonic of the stimulation frequency 4.8 Hz, showing prominent auditory networks and cingulate gyrus in PL and DMN in RS. Given the harmonic relationship and significant overlap with fundamental auditory networks, we interpret the attunement of the 4.8 Hz network as being functionally related to auditory processing.

In the alpha range (8-13 Hz), the spatial activation patterns displayed a relevant reorganization across conditions. In RS, the maximally attuned network at 10.9 Hz showed a spatial distribution of the first two GED components centered in the parieto-occipital regions, while higher alpha frequencies showed a strong recruitment of sensorimotor regions. In PL, the alpha peak shifted to 12.1 Hz, and sensorimotor regions replaced parieto-occipital regions across the entire alpha range. Overall, the rearrangement of alpha networks consisted of a focal cluster of neighboring regions shifting from parieto-occipital areas at RS to sensorimotor regions during PL. Interestingly, **Figure 2** demonstrates a posterior-to-anterior spatial gradient in the alpha frequency range, which shifts as frequency increases. While this gradient is present in both conditions, indicating that these are inherent brain dynamics, the significant change in eigenvalues suggests that attunement to a preferred peak frequency is condition dependent. This spatial reorganization is qualitatively different from the transitions away from the DMN observed at lower frequencies, where the emergence of frequency-resolved networks over aperiodic background activity was linked to a focal convergence from a broader spatial distribution.

In the beta range (13 – 30 Hz), the spatial activation patterns were consistent across both RS and PL, pointing to a primarily involvement of sensorimotor regions for the first two components in both experimental conditions. In the gamma range (30 – 97.6 Hz), similar to beta, the spatial activation patterns remained consistent across conditions. Here, gamma activity was broadly distributed, encompassing bilateral insula, inferior temporal cortex, hippocampal regions, frontal operculum and inferior frontal gyrus. The topographies of the gamma networks are shown in **Figure 3** and **Figure S1**.

**Figure 3:**
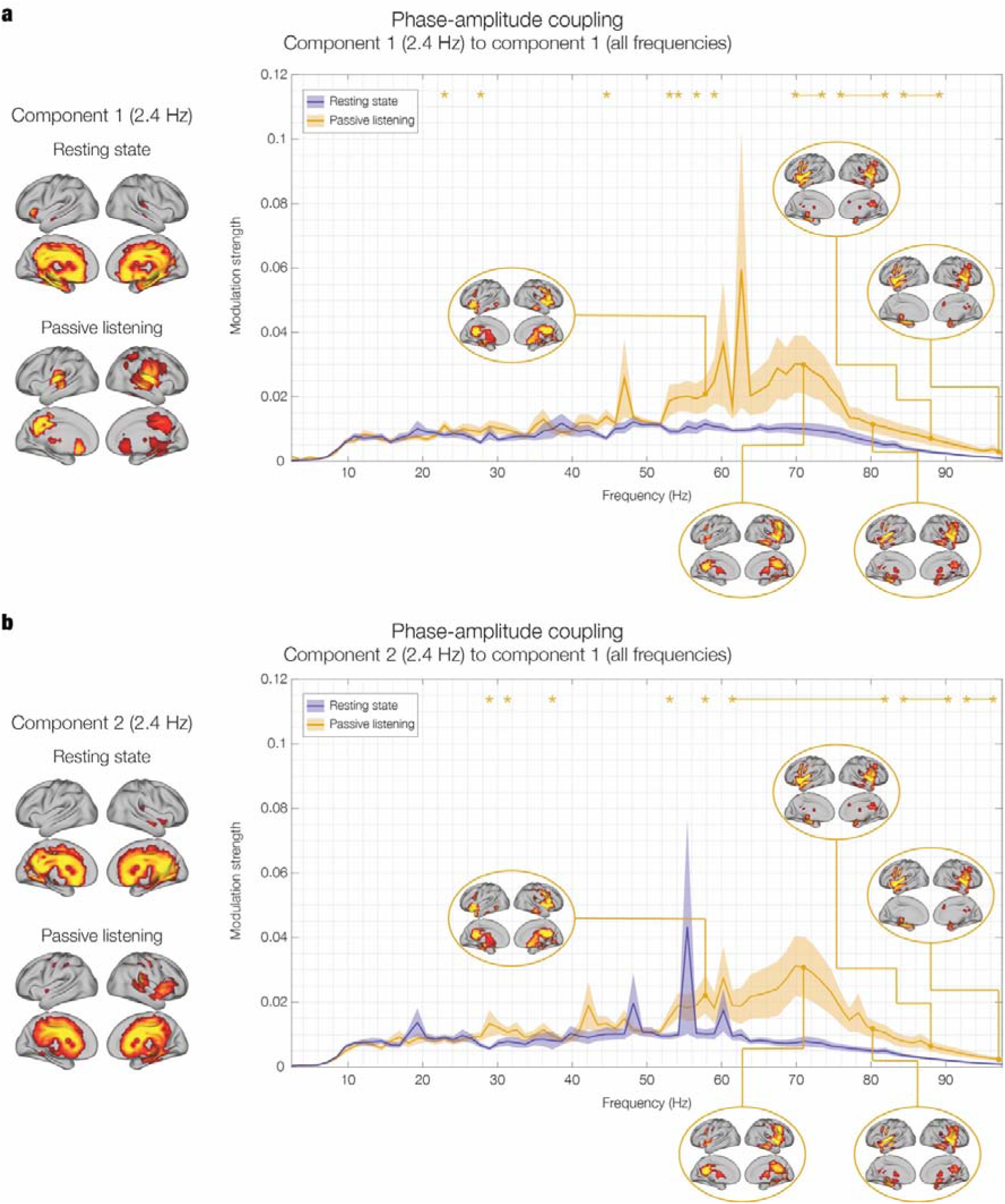
Network landscape: cross-frequency coupling during rest and passive listening. The figure shows the interaction between each of the two low-frequency 2.4 Hz auditory networks and all the higher-frequency networks during resting state (R) and passive listening (PL). The plots display the modulation strength, a measure of how the phase of the low-frequency network component modulates the amplitude of the other network components. The solid lines represent the mean modulation strength averaged across 26 participants (N = 26), with shaded areas indicating the standard error of the mean (SEM). Stars denote frequency bins where significant differences were detected using a right-tailed signed-rank Wilcoxon test, following FDR correction for multiple comparisons. **a - Component #1 (2.4 Hz)**: The left panel shows the spatial activation patterns for component #1 at 2.4 Hz in both resting state and listening conditions. In R, the activation is concentrated in medial temporal regions, including the hippocampus, while during listening, it shifts towards primary auditory areas, particularly Heschl’s gyrus, indicating the auditory network’s engagement with the rhythmic stimulus. The right panel displays the PAC results, showing significant modulation of high gamma band activity (70-90 Hz) by the 2.4 Hz component during listening. This modulation is notably stronger than in the resting state, suggesting enhanced cross-frequency coupling driven by auditory processing. **b - Component #2 (2.4 Hz)**: The left panel shows the spatial activation patterns for component #2 at 2.4 Hz, which involve a broader network during both conditions, including auditory and medial temporal regions. In the listening condition, this component also recruits additional prefrontal regions, indicating a higher-level integration of auditory information. The right panel shows that, similar to component #1, the modulation of gamma band activity by component #2 is significantly stronger during listening, particularly in the 60-80 Hz range, reinforcing the finding that auditory stimulation enhances the coupling between low-frequency auditory rhythms and high-frequency gamma oscillations. These results highlight the dynamic reconfiguration of cross-frequency interactions in response to auditory stimulation, with a clear enhancement of PAC between low-frequency auditory components and high-frequency gamma networks. The enhanced modulation of gamma band activity by the low-frequency auditory networks during listening suggests heightened cross-frequency coupling, indicative of increased interaction across neural networks in response to external stimuli. This underscores that gamma oscillations play a key role in processing rhythmic auditory stimuli, mediated by the interaction with auditory networks attuned to the stimulation frequency.

To ensure the robustness of our methodology against sample variability and changes in MEG processing, we conducted a series of replication analyses using the same pipeline. **Figure S2** demonstrates that the RS network landscape was successfully replicated in two larger, independent resting-state datasets, one of which included participants within a very narrow age range to further reduce individual variability. **Figure S3** illustrates the replication of the RS and BL network landscapes using shorter time durations, showing that just 30 seconds of recording are sufficient to reproduce our findings. **Figure S4** presents an exact replication of the results on the current dataset, reconstructing anatomical sources using only gradiometers instead of magnetometers. This confirms that our method is invariant to the choice of sensors for beamforming implementation (see ^53^, for a systematic comparison of different implementations). Further details and discussion of these replication results, supporting the findings reported in the main manuscript, are provided in the captions of the respective supplementary figures. Detailed statistical results for all supplementary analyses are reported in **Table S4**.

### Cross-frequency coupling

To investigate the dynamics of interaction across the separated networks, we analyzed the cross-frequency coupling (CFC) between selected combinations of their activation time series. Specifically, we assessed whether activation in networks across all frequencies was modulated by the auditory networks attuned to the stimulation frequency. To do so, we calculated the power distribution of all frequency-resolved components over the phase time series ^54–56^ of the first component of the 2.4 Hz modulator. Given the responses elicited in the secondary auditory network, we repeated the procedure for the second component of the 2.4 Hz modulator. Motivated by the empirical observation that the power distributions are well approximated by a sinusoidal function, we fitted a sinewave to each distribution and used its amplitude as an index of modulation strength ^34^.

As illustrated in **Figure 3**, we tested the hypothesis that PL would result in a significantly stronger modulation than RS over the networks landscape. We performed a Wilcoxon signed-rank test on the modulation strength of PL vs RS, over the entire frequency spectrum and independently for the first two 2.4 Hz GED components. The results were corrected for multiple comparisons using FDR.

The statistical tests returned the following main significant frequency windows for the first 2.4 Hz component (**Figure 3a**): 71.0 – 73.5 Hz; 77.0 – 81.9 Hz; 84.3 – 87.9 Hz (p < 0.01, false-discovery rate FDR-corrected; for an extensive report of the exact statistics, see **Table S2**). **Figure 3b** shows the results of the same test for the modulation by the second 2.4 Hz component, which resulted in the following main significant frequency windows: 63.8 – 81.9 Hz; 84.3 – 89.1 Hz; 92.7 – 96.3 Hz (p < 0.02, false-discovery rate FDR-corrected; for an extensive report of the exact statistics, see **Table S2**).

These results indicate that cross-frequency coupling was significantly stronger in PL compared to RS, selectively for the gamma band. Specifically, the phase fluctuations of oscillations at 2.4 Hz modulated the power of higher-frequency components to a larger extent during auditory stimulation, suggesting a heightened state of interaction across networks driven by the external rhythmic stimulus. This level of analysis provides further insight into the interplay between different neural oscillations during auditory processing and how it relates to the configuration of different networks. Importantly, it showed that the role of the gamma network in sensory processing is mediated by the coupling with primary auditory networks rather than by an internal frequency attunement.

Detailed statistical results for the modulation of all components over the frequency spectrum are reported in **Table S2**.

### Randomizations

In order to systematically test the principle of dissociation between the temporal and spatial dimensions of FREQ-NESS, we performed two types of randomizations on our RS data in voxel space, and iterated GED on the same data in the four shuffled conditions. This was done only in RS since it was superfluous to conduct the same procedure in both experimental conditions. The randomization strategies were carried out as follows, operating on a matrix of brain voxels with two dimensions (space and time): (1) *label randomization*: the brain voxel indexes were shuffled, in a consistent manner over the entire time series (i.e. no changes were made to the time series but the time series were entirely reassigned to different brain voxels); (2) *point-wise label randomization*: the voxel indexes were shuffled row-wise, independently for each timepoint.

By applying these randomization techniques, we sought to disrupt the temporal and spatial structures of the data independently. **Figure 4** shows the effects of the randomization strategies on the network landscape in RS. Label randomization (1) disrupted the spatial activation patterns without altering the eigenspectrum. This preserved the frequency content of the time series and the variance explained by the GED components but returned physiologically meaningless activation patterns. Point-wise label randomization (2) disrupted both the spatial activation patterns and the eigenspectrum, indicating that reshuffling the voxels for every time-point disrupted the temporal structure of the multivariate signal and, therefore, the frequency content of the estimated networks. Here, the variance explained by the first GED components decreased to an average of 0.06% across frequencies, flattening the exponential decrease in the variance explained by progressive GED components at each frequency. The latter randomization provides insights into the distribution of variance explained by GED components when attributable to chance.

**Figure 4:**
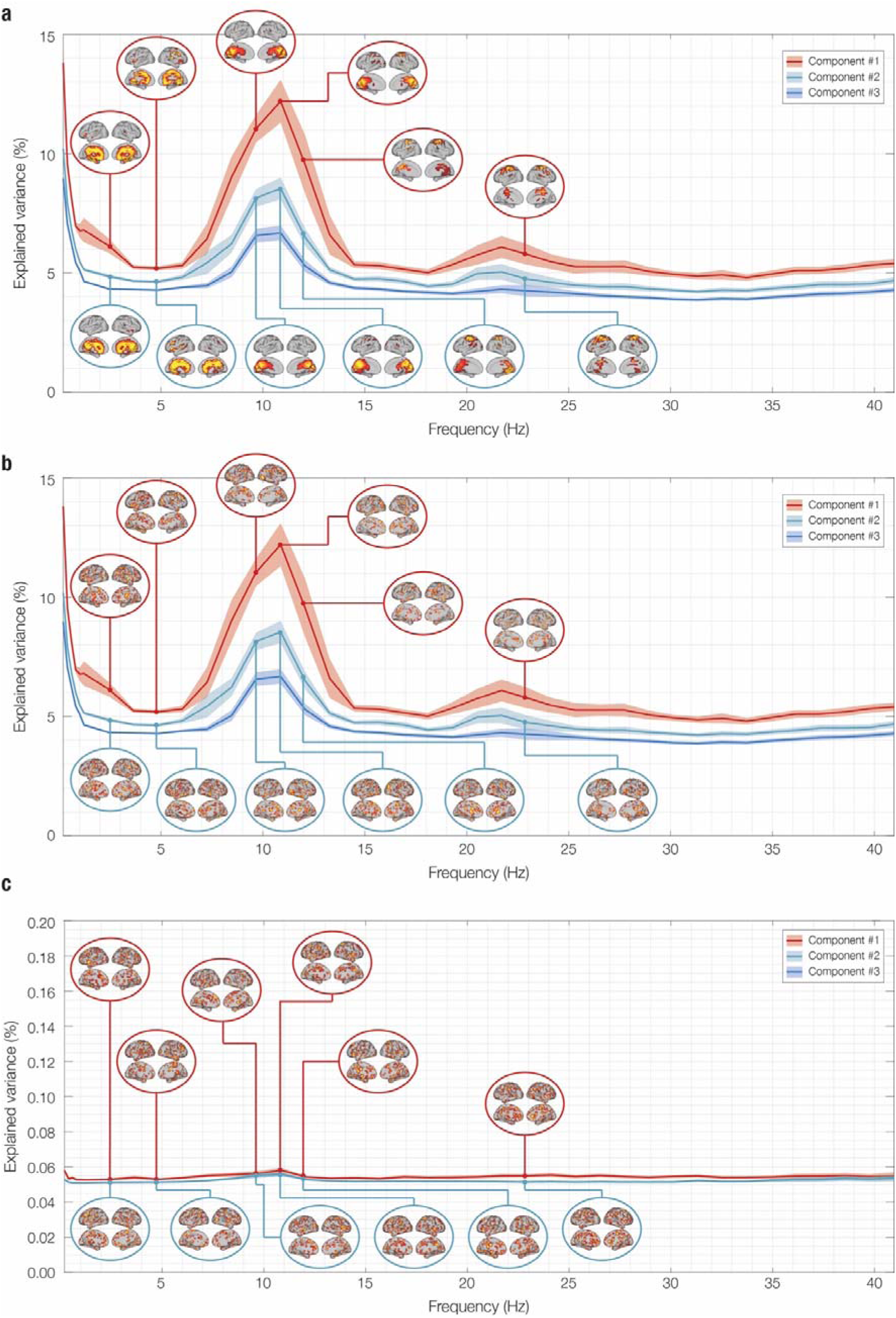
Network landscape of randomized resting state data: eigenspectrum and spatial activation patterns. This figure illustrates the frequency-resolved brain networks obtained using FREQ-NESS applied to the original data, randomized with two different strategies. 1)Label randomization: The brain voxel indices were shuffled consistently across the entire time series, meaning the temporal data remained intact but was reassigned to different brain voxels; 2) Point-wise label randomization: The voxel indices were shuffled independently for each time point, disrupting the temporal structure. As shown in Figure 2, first element of the network landscape is the eigenspectrum, which shows the eigenvalues representing the percentage of variance explained by the GED components across frequencies. Solid lines denote the mean normalized eigenvalues across participants, with shaded areas indicating the standard error of the mean (SEM). The second element depicts spatial activation patterns associated with the top GED components at frequencies with the highest explained variance. Each pattern illustrates the contribution of each brain voxel to the corresponding network. For enhanced visualization, the eigenspectra are presented for the first three GED components (sorted by explained variance) across a frequency range of 0.2 to 40.9 Hz. The spatial topographies of the first two components are shown for frequencies with the highest explained variance. **a – Original data**: To allow an easier comparison, we report the same results for resting state data shown in Figure 2. **b – Randomization 1 (Label)** preserved the eigenspectrum but disrupted the spatial activation patterns, maintaining the frequency content and variance explained by GED components but yielding physiologically meaningless patterns. **c – Randomization 2 (Point- wise label)** disrupted both the eigenspectrum and spatial activation patterns, flattening the exponential decrease in variance explained by subsequent GED components. This suggests that reshuffling voxels for each time point disrupts the temporal structure of the signal and reduces the variance explained by GED components to approximately 0.06% across frequencies, highlighting potential variance contributions attributable to chance. The solid lines represent the mean normalized eigenvalues averaged across 26 participants (N = 26), with shaded areas indicating the standard error of the mean (SEM). A two-tailed signed-rank Wilcoxon test was conducted separately for each randomization against the original data. Since Randomization 1 yielded no significant differences, while Randomization 2 showed significant differences across all frequency bins following FDR correction for multiple comparisons, statistical significance is not reported in the figure to avoid redundancy.

To statistically assess changes in the RS network landscape resulting from the randomization strategies, we performed a series of Wilcoxon signed-rank tests. Specifically, we compared the original RS data (see **Figure 4a**) with Randomization 1 (Label, see **Figure 4b**) and Randomization 2 (Point-wise label, see **Figure 4c**) across the entire frequency spectrum for the top three eigenvalues. FDR correction was applied to account for multiple comparisons. As expected, Randomization 1 showed no significant differences, while Randomization 2 yielded significant differences across all frequency bins (p < 0.001, FDR-corrected). These results statistically confirm our qualitative observations based on visual inspection.

## Discussion

Scientific research has long established that the brain is a complex, dynamic system organized into networks ^1,2,27,28,44,57^. However, much of the existing evidence has arisen from studies that narrowly focused on predefined brain regions or canonical frequency bands. Additionally, the use of complex methods often hindered straightforward interpretations. While these approaches have yielded valuable findings, they also present challenges in integrating the brain’s temporal and spatial dynamics into a cohesive framework that is based on minimal assumptions and allows for clear interpretations. To address these challenges, we developed a novel analytical pipeline named *FREQuency-resolved Network Estimation via Source Separation* (FREQ-NESS) and applied it to MEG recordings during both resting state and continuous auditory rhythmic stimulation. This flexible, data-driven approach is based on three main assumptions supported by previous evidence: (1) a brain network exists when independent brain voxels covary over time ^58–60^, (2) during both resting state and tasks involving external rhythmic stimulation, brain networks operate at different frequency bands ^11,30^, and (3) the GED components generated by FREQ-NESS are directly interpretable as brain networks. Here, the variance explained by these components indicates the relative significance of the networks for each frequency band and experimental condition, while the associated activation patterns and time series provide insights into the spatial and temporal extent of the networks.

### Frequency-resolved brain networks during resting state and passive listening

As expected from successful source separation ^11,33^, we observed a pronounced exponential decay in the variance explained by the GED components at key frequencies (see **Figure S1** for an overview across all frequencies and **Figure S5** for a detailed view of the key frequencies). With the eigenvalues sorted in descending order of explained variance, this pattern indicated that the top components account for disproportionately higher amounts of variance. Analyzing additional components would result in exponentially diminishing returns, increasing the risk of multiple comparisons with minimal benefit and reduced parsimony. Notably, for frequencies where the decay was more pronounced, only the first three eigenvalues stood out prominently in the distribution. Based on this observation, we focused our analysis on the most prominent networks associated with these eigenvalues to reduce data dimensionality while minimizing information loss (see **Table S3** for cumulative explained variance by the top components).

In resting state, the variance explained by these GED components showed a 1/f decay from the slow to high frequencies in the delta range, a prominent alpha peak at 10.9 Hz, and a reduced yet relevant beta peak at 22.9 Hz (see **Figure 2a**). During listening, we instead observed an outstanding peak at the stimulation frequency of 2.4 Hz and its first harmonic 4.8 Hz, along with a prominent alpha peak at 12.1 Hz, and beta peak at 22.9 Hz (see **Figure 2b**).

In this study, the 2.4 Hz frequency within the delta range was of particular interest because it matched the adopted stimulation frequency. During resting state, the variance explained by the GED components exhibited an exponential decay with increasing frequency, with no prominent peaks at 2.4 Hz. However, the spatial activation patterns across all delta band components, including at 2.4 Hz, were highly comparable and displayed the characteristic configuration of the DMN (**Figure 2**). The DMN is widely recognized as the dominant activation mode of the brain at rest, primarily involved in self-referential and introspective activities rather than external stimulus processing (for a review, see ^61^). Notably, no prominent eigenvalue associated with this network emerged from the spectrum, indicating that the DMN exhibits low frequency-specificity according to our criteria for source separation. This finding suggests that the DMN’s spatial activation pattern persists across different frequency bins within the delta band when the brain is not engaged in a specific task, underscoring its role in brain activity at rest ^62^.

Beyond resting state, when GED was attuned to 2.4 Hz in the listening condition, the eigenspectrum was dominated by one eigenvalue explaining 18% and a second eigenvalue explaining 7% of the total variance (see **Figure 2b**). This finding indicates that two components of the brain voxel signal maximally attuned to the stimulation frequency, based on their narrow spectral density. Inspection of their spatial projections onto voxel space revealed focal patterns in auditory areas, providing a clear physiological explanation for the frequency attunement of brain activity to the stimulation. To elaborate, we inferred that two brain networks were recruited to process the rhythmic stimuli, coherent with low- and higher-level processing of auditory stimuli previously reported in the literature ^50,63–66^. The focal activation in the Heschl’s gyrus observed in the first network, with asymmetry favoring the right hemisphere previously reported in healthy adults ^64,67^, is associated with early components of auditory processing ^50,63,68^. The activation of the second network appeared more widespread, prominently involving medial temporal regions such as hippocampal regions, insula, inferior temporal cortex as well as cingulate gyrus and ventro-medial prefrontal cortex, which were previously connected to later stages of auditory processing ^52,69^.

Although the second network partially overlaps with the DMN on a spatial level, its narrow attunement suggests its genuine involvement in auditory processing rather than being a residual of the DMN operating in the background.

The second frequency identified in our analysis falls within the alpha range, which is ubiquitous in the brain and therefore prominent in most M/EEG sensors ^70^. Interestingly, the peak and distribution of the variance explained by the top alpha networks differed across resting state and passive listening. At rest, it approximated a symmetric Gaussian curve peaking at 10.9 Hz. During listening, the left side of the curve flattened, and the peak shifted higher to 12.1 Hz. During resting state, the spatial activation pattern of the peak components displayed the stereotypical parieto-occipital activation characteristic of the brain at rest ^20,21,71,72^. In contrast, during listening, the focal activation was centered in the sensorimotor areas, highlighting the involvement of alpha mu-rhythms typically observed in these regions^72–74^.

Notably, a more fine-grained inspection of the left- and right-sides of the eigenvalues’ distribution in the alpha range revealed that even during resting state, the focus of the spatial extent of the alpha network transitions anteriorly towards the pre-central gyrus and sensorimotor regions with increasing frequency (see **Figure 2**). Indeed, while not as dominant in the distribution, higher alpha is still spatially concentrated in sensorimotor areas such as supplementary motor area (SMA). What occurred during listening is that the sensorimotor alpha network at 12.1 Hz became more prominent relative to the same network at rest. We interpret this spatial rearrangement of the alpha network as underpinning readiness for action, given the natural human inclination to synchronize with periodic auditory stimuli ^75–77^. This is particularly true for the involvement of the SMA, which is thought to inform expectations of auditory periodic stimuli by representing cyclic sensorimotor processes, enabling sensory predictions in absence of movement and ultimately overt behavioral synchronization with a beat ^78^.

The third frequency identified in our analysis was found in the beta range, peaking at 22.9 Hz and exhibiting the same spatial activation patterns in both conditions. The focal activation over the precentral gyrus aligns with the expected topography at rest, as beta is a dominant spectral mode of motor cortices ^20,72,73^. Interestingly, the same pattern was observed during the listening condition, where different outcomes might have been anticipated. For example, given the contribution of motor beta dynamics to auditory processing ^79–81^, one might reasonably expect either a significant change in the explained variance of this motor network or the emergence of an additional auditory network dominated by beta activity ^81–83^. At the same time, beta oscillations in sound and music perception are associated with processes such as internal representation, prediction, and hierarchical subdivision of sounds, making it sensitive to the task and complexity of the stimuli ^81,83–85^. In light of our results and given the extreme simplicity of the isochronous tone sequences presented to our participants, we conclude that the emergence of a prominent beta network was likely unnecessary to process the stimulation in our experimental condition. Furthermore, fluctuations in beta power induced by rhythmic stimuli are dissociable by the overall level of beta power ^34^, which may have left the variance explained by the beta network unchanged across conditions.

### FREQ-NESS as an optimized analytical pipeline for brain network estimation

FREQ-NESS captured a dissociation between temporal and spatial dimensions in brain activity, successfully identifying the classical networks commonly associated with the resting state, such as the default mode network, alpha-band network, and motor-beta network ^20,21,44,86^. Notably, FREQ-NESS makes advances beyond other methods by providing a comprehensive overview of simultaneous brain networks at rest and revealing three scenarios resulting from auditory stimulation compared to the resting state baseline: (1) emergence of prominent networks attuned to the stimulation frequency (e.g., transition from DMN to auditory networks attuned to the stimulation frequency); (2) topographical rearrangement of endogenous oscillations already prominent in the brain activity at rest (e.g., rearrangement of alpha networks); (3) maintenance of the status quo for networks operating at frequencies not relevant to stimulus processing (e.g., invariance of the beta motor network).

While FREQ-NESS represents a novel analytical pipeline for studying brain networks, it is grounded in well-established concepts and analytical methods commonly used in neuroscience ^11,23,24,31,40^. Indeed, over the last decade, GED has been applied to sensor data matrices measured by M/EEG with the explicit purpose of separating latent processes in the brain projecting and mixing to the recording at the scalp level (e.g., ^26,33–36,40–43)^. However, a major limitation of such approaches arises from the fact that brain dipoles are postulated to be inaccessible latent sources, and therefore the physiological interpretability of spatial activation patterns is limited to a model of diffusion across sensors due to volume conduction (for an extensive discussion on the interpretability of the eigenvectors, see ^58,85–87)^. One major insight behind FREQ-NESS is that applying GED on a data matrix of reconstructed brain voxels enables a direct and physiologically meaningful estimation of brain networks. In addition to GED, other methods have been previously explored for separating sources and deriving insights into brain networks. For instance, some studies gathered insights about network configurations in the brain by using Independent Component Analysis (ICA) ^60,87–89^ and Principal Component Analysis (PCA) ^59,90–92^, especially in relation to fMRI data. However, while valuable, the signal measured by fMRI has very low temporal resolution on the order of seconds and does not directly capture the neurophysiological signal produced by the brain. This means it lacks the ability to capture relevant frequency information and cannot make inferences on the fast-scale neurophysiological processes occurring in the brain at rest and during stimulus processing. As a result, when applied to fMRI data, ICA and PCA cannot identify frequency-resolved brain networks using detailed, temporally rich neurophysiological signals.

In neurophysiology, the use of linear decompositions to derive brain networks is relatively rare. An interesting example is the study by Brooks et al. ^93^, where ICA was applied to MEG data across canonical broad frequency ranges, resulting in network configurations comparable to those observed in fMRI during resting state ^93^. While valuable, ICA is not specifically designed to separate frequency-resolved networks, lacking the flexibility to define criteria for source separation, such as contrasting broadband and narrowband frequencies. Additionally, ICA does not provide information about the variance explained by the components, making it challenging to select the most relevant ones. Moreover, the study did not employ a systematic frequency-resolved approach. Building on this, to explore an alternative and computationally simpler method, in the current study we also implemented a variation of the FREQ-NESS analytical pipeline using PCA on narrowband-filtered brain voxel data. While PCA does not explicitly contrast narrowband and broadband covariance structures, it offers a straightforward eigendecomposition framework within the same analytical paradigm. A detailed description of this approach, along with comparative results against GED, is provided in *Supplementary Information* (**Figure S6**). Our findings indicate that while PCA produces network landscapes with some qualitative similarities, it exhibits reduced spectral precision and significantly lower spatial specificity. Furthermore, PCA introduces systematic biases due to its variance-maximizing nature, rather than optimizing for frequency specificity. This results in a persistent broadband 1/f component blending with the eigenvalue distribution across frequencies, as well as a background DMN configuration merging with expected activation foci at specific frequencies. These results show that narrowband filtering alone does not ensure de-mixing, as PCA’s constraints cause networks to remain entangled in time and space. Overall, our findings reinforce the superiority of GED for isolating oscillatory activity in multivariate neurophysiological data while recognizing PCA as a viable, albeit suboptimal, alternative for frequency-specific network decomposition. In contrast, we have presented evidence that FREQ-NESS implemented with GED is an effective solution to separate brain networks in a data-driven manner based on their frequency-specificity, under three fundamental assumptions which widely align with existing research. The first assumption is that the multivariate time series characterizing the brain voxels is a linear combination of sources mixed with noise, and that a brain network exists when independent brain voxels covary over time. This allows the estimation of brain networks via eigendecomposition of the covariance matrices computed on the broadband and frequency-resolved voxel data. The second assumption is that oscillatory dynamics are the dynamics of interest, whereas broadband aperiodic components of brain activity are technically treated as noise to enhance contrast with a reference covariance matrix ^94^. This assumption constrains the solution space while giving the researcher control over the criteria for separating the sources, making GED an ideal tool to perform frequency-resolved blind source separation ^93^ (but see ^95^, for other approaches accounting for aperiodic components of brain activity). Furthermore, this second assumption is also backed by substantial evidence that preferred frequency modes characterize cognitive processes ^96^ and dominate the activity of specific brain regions ^23^. The third assumption is that the GED components generated by FREQ-NESS are directly interpretable as brain networks. Here, the variance explained by these components indicates the relative prominence of the networks for each frequency band and experimental condition, while the associated activation patterns and time series provide insights into the spatial and temporal extent of the networks ^2,25,95,96^.

### Cross-frequency coupling interaction between brain networks

FREQ-NESS not only delivers insights into brain network estimation but also offers flexibility for integration with other analytical methods, enhancing their capabilities. For example, FREQ-NESS enables the study of cross-frequency coupling (CFC) between entire brain networks, rather than between individual pairs of brain regions. More specifically, computing CFC between the narrowband filtered time series of different brain networks enables the quantification of their interactions in the form of phase-to-amplitude coupling ^2,25,97,98^. To test the hypothesis that high-frequency network activity is modulated by a low-frequency oscillation attuned to the stimulation frequency (2.4 Hz), we computed an index of modulation based on the distribution of high-frequency power values over low-frequency phase ^34,54–56^ . Our results revealed that power in gamma networks between 63.8 Hz and 96.3 Hz is significantly modulated by low-frequency oscillators in the two main auditory networks attuned to the 2.4 Hz stimulation frequency. This result is coherent with the established notion that oscillations in the gamma band are induced by the presentation of repetitive auditory stimuli and are associated with their bottom-up processing ^82,99–102^. Notably, our study did not highlight a specific role for the auditory cortex in the gamma network dynamics, which instead primarily recruited insula, inferior temporal cortex, hippocampal regions, frontal operculum and inferior frontal gyrus. This suggests that modulated gamma activity may not originate directly in the auditory cortex but rather be mediated by interactions within low-frequency auditory networks.

### Conclusions and future perspectives

This work presents several implications and potential future perspectives. A key implication is that FREQ-NESS is designed for use with continuous MEG recordings co-registered with MRI data. This tool is valuable for researchers who have a specific hypothesis about activity at a particular frequency of interest and for those who aim to explore the entire network landscape without the a priori assumptions previously discussed. Consequently, FREQ-NESS can be particularly useful in analyzing datasets involving continuous stimuli such as music or speech to uncover neural responses and connectivity patterns, where the temporal structure of the stimulus can be decomposed in frequency components and mapped to neurophysiology^6,37,39^ and domain-specific networks are characterized by spectral fingerprints ^85,101–104^.

In absence of stimulation, other datasets of interest include the investigation of states of consciousness, involving continuous resting state recording carried out in conditions of meditation or psychedelics, where altered states of consciousness have been associated with changes in the power spectral density over the whole brain ^87,103–106^. Along this line, large-scale comparisons of resting state recordings across populations in a between-subject design would enable to establish a normative landscape of frequency-resolved networks at rest and, eventually, during tasks.

FREQ-NESS demonstrated high reliability even when analyzing data from individual participants, as evidenced by the low standard errors in our statistical measures. This reliability, further supported by extensive replications across independent datasets and shorter recording fragments (see *Supplementary Information*), enables FREQ-NESS to produce robust results from small sample sizes, facilitating the generation of individualized data. Such data could be crucial for clinical applications that aim to diagnose pathological changes in the frequency-specific brain network patterns of clinical populations compared to healthy individuals.

Another future direction consists of deploying tasks to elicit dynamics in networks of interest. One concrete example comes from perturbation paradigms in the context of sensorimotor synchronization. As previously pointed out ^33^, the network estimation at the level of the anatomical sources could potentially solve the long-standing problem of neural entrainment (see ^65^), disentangling the different networks underlying endogenous oscillations and periodic evoked responses within critical time windows of frequency alignment in response to tempo changes ^33^.

Our study showed that GED is a robust and principled approach for estimating distinct frequency-specific brain networks. However, future research could expand on our work by not only comparing GED and PCA but also exploring other methods. While these alternatives may not be ideal for our specific goals, techniques like Multivariate Empirical Mode Decomposition (MEMD) ^22^ could offer valuable insights and fresh perspectives.

Finally, it is worth noting that our analytical approach normalized the beamforming weights during source reconstruction to mitigate issues such as bias towards the center of the head in the reconstructed data. However, in this study, we did not record empty-room data, which could have provided a valuable comparison between actual data and the general noise level in the environment. Future studies may consider incorporating this step to further evaluate the robustness of our methodology.

To conclude, the current study demonstrated the effectiveness of FREQ-NESS as a multivariate analytical pipeline to provide a comprehensive and physiologically interpretable perspective on the brain functioning, in terms of a landscape of frequency-resolved brain networks.

## Methods

### Participants

A total of 29 participants (N = 29) took part in the study (17 females, 12 males). The age of the participants ranged from 19 to 50 years (mean age = 26.45 ± 7.37 years). The group consisted of four first-year music students and 25 non-musicians. Except for one participant in the latter group, all participants had less than two years of formal musical theory training (mean age = 0.23 ± 0.47 years) and less than five years of formal musical instrument training (mean years = 0.59, ± 1.10). None of the participants have been diagnosed with any neurological or psychiatric disease, nor had hearing impairment. All participants except one were right-handed. Of the 14 countries represented by our sample, Danish was the largest national group (24%). Participants were compensated for their time with vouchers for online shopping. To be noted, two participants did not undergo the passive listening (PL) recording, and one participant was discarded due to technical issues during the data collection, leaving us with twenty-six (n = 26) participants eligible for within-subject comparisons in both conditions.

### Experimental procedure

Upon arrival, participants were instructed about the experimental procedures before signing the informed consent form. They were given the possibility to withdraw their participation in the study at any time. A researcher was always present to provide assistance or clarifications when needed. Participants underwent MEG recordings under two conditions. In the PL condition, participants listened to rhythmic auditory stimulation while watching a silent movie. The stimuli consisted of an isochronous sequence of 300 Hz sine tones of 100 ms duration, with 10 ms fade-in and 10 ms fade-out, presented at a rate of 2.4 Hz (400 ms inter-onset interval) for a total duration of five minutes. Prior to the measurements, the sound intensity was set at 50 dB above the minimum hearing threshold of each participant. In the resting state RS condition, no auditory stimulation was presented while participants watched the same silent movie for a total duration of 10 minutes. Only the first five minutes of the resting state recording were analyzed, in order to match the passive listening condition. The present study was conducted on a subset of a larger dataset acquired at Aarhus University and approved by the Ethics Committee of the Central Denmark Region (De Videnskabsetiske Komiteer for Region Midtjylland) (ref. 1-10-72-411-17). The experimental procedure was carried out according to the Declaration of Helsinki.

### Data Acquisition

The MEG data were collected in a magnetically shielded room at Aarhus University Hospital (AUH) in Aarhus, Denmark, using an Elekta Neuromag TRIUX MEG scanner with 306 channels (Elekta Neuromag, Helsinki, Finland). The recordings were sampled at 1000 Hz with analog filtering set between 0.1 and 330 Hz. Prior to recording, the participants’ head shapes and the positions of four Head Position Indicator (HPI) coils were registered relative to three anatomical landmarks using a 3D digitizer (Polhemus Fastrak, Colchester, VT, USA). This information was later used to align the MEG data with MRI anatomical scans in co-registration. During the MEG sessions, the HPI coils continuously tracked head position for movement correction during pre-processing. Additionally, two sets of bipolar electrodes recorded cardiac rhythm and eye movements, which facilitated the removal of electrocardiography (ECG) and electrooculography (EOG) artifacts during pre-processing.

MRI scans were obtained using a CE-approved 3T Siemens MRI scanner at AUH, with structural T1 (mprage with fat saturation) data recorded at a spatial resolution of 1.0 x 1.0 x 1.0 mm. The sequence parameters were set as follows: echo time (TE) = 2.61 ms; repetition time (TR) = 2300 ms; reconstructed matrix size = 256 x 256; echo spacing = 7.6 ms; bandwidth = 290 Hz/Px. The MEG and MRI recordings were conducted on separate days.

### MEG Data Pre-Processing

The raw MEG sensor data, consisting of 204 planar gradiometers and 102 magnetometers, were initially pre-processed using MaxFilter 69 ^107^ to reduce external interferences. Signal space separation was applied with the following MaxFilter parameters: spatiotemporal signal space separation (SSS); down-sampling from 1000 Hz to 250 Hz; movement compensation with cHPI coils (default step size: 10 ms); and correlation limit of 0.98 between inner and outer subspaces to reject intersecting signals during spatiotemporal SSS. As commonly practiced in the literature ^108,109^, data were downsampled by a factor of four to facilitate the computational feasibility of our analyses without significant data loss. Frequencies up to 97.6 Hz were explored in our analysis, a boundary which is still well below the Nyquist frequency of 125Hz given a sampling rate of 250 Hz. The data were then converted to Statistical Parametric Mapping (SPM) format for further pre-processing and analysis in MATLAB (MathWorks, Natick, MA, USA) using custom-built codes (LBPD, https://github.com/leonardob92/LBPD-1.0.git) and the Oxford Centre for Human Brain Activity (OHBA) Software Library (OSL) ^110^, which builds on Fieldtrip ^111^, FSL ^112^, and SPM ^113^ toolboxes. The continuous MEG data were visually inspected using the OSLview tool to identify and remove the artifacts. Independent component analysis (ICA) (OSL implementation) was used to eliminate eye-blink and heart-beat artifacts from the brain data^114^. The original signal was decomposed into independent components (ICs), and then each IC was correlated with activity recorded by the EOG and ECG channels. ICs showing a correlation at least three times higher than the others with either the EOG or the ECG were labelled for removal. To ensure the accuracy of this procedure, we also visually inspected the activation time series and the topographic scalp distribution of the labelled ICs, to confirm the expected stereotyped patterns generated by eyeblinks and heartbeats. When both the correlations and visual inspections indicated that an ICA component was due to eyeblink or heartbeat activity, those components were discarded. The signal was then reconstructed by back projecting the remaining components in the MEG sensor space. The continuous recordings of each experimental condition were segmented into a single 5-minute epoch.

### Anatomical Source Reconstruction

MEG is highly effective in detecting whole-brain activity with excellent temporal resolution. However, in order to provide a physiologically interpretable characterization of the brain networks, it was crucial to estimate the anatomical sources projecting their activation at the scalp level. Since MEG records neural signals outside the head without directly indicating their specific brain sources, these have to be reconstructed by tackling an inverse problem. For this purpose, we used the established beamforming method ^115–117^ (**Figure 1b**), which integrates custom-built code with routines from OSL, SPM, and FieldTrip.

Beamforming algorithms consist of two main steps: (1) creating a forward model for the projection of the brain sources to the sensors and (2) computing the inverse solution. The forward model treats each brain source as an active dipole (brain voxel) and predicts how the unitary strength of each dipole is reflected across all MEG sensors. Here, we used only the magnetometer channels with an 8-mm grid, yielding 3559 dipole locations (voxels) throughout the brain. This choice was based on magnetometers’ better detection of deep brain activity compared to gradiometers, which are more sensitive to cortical activity near each gradiometer ^108,109^. After co-registering individual structural T1 data with fiducial points (head landmarks), we computed the forward model using the widely adopted “Single Shell” method detailed by Nolte 83. The result, called the “leadfield model,” was stored in matrix L (sources x MEG channels). For the participants lacking structural T1 data, we used a template (MNI152-T1 with 8-mm spatial resolution) for leadfield computation. Next, we computed the inverse solution using beamforming. This process sequentially applies different weights to source locations to isolate each source’s contribution to the MEG signal at every time point. The beamforming inverse solution involves several steps. The MEG sensor data (B) at time *t* is described by the equation:

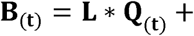

where *L* is the leadfield model, *Q* is the dipole matrix containing each dipole’s activity over time, and L represents noise (see ^117^ for details). To solve for *Q*, the weights (*W*) must be computed, and then applied to the MEG sensors at each time point:

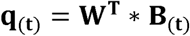

To obtain *q*, the weights *W* are calculated as shown in equation:

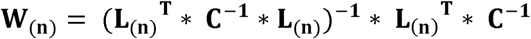

where *I* is the covariance matrix between MEG sensors, calculated from the single 5-minute epoch. Leadfield model computation was performed for three main orientations of each brain source (dipole) according to Nolte ^118^, but orientations were reduced to one using the singular value decomposition algorithm to simplify the beamforming output ^119,120^.

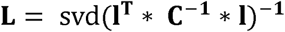

Here, *l* represents the leadfield model with three orientations, while *L* is the simplified one-orientation model used in the equation above. The weights were applied to each brain source and time point, with the covariance matrix computed from the continuous signal estimated by concatenating trials of all experimental conditions. The weights were normalized following the method described by Luckhoo et al. ^121^ to mitigate reconstruction bias toward the center of the head, a critical step, particularly when empty-room data are unavailable. Weights were applied to neural activity from the single epoch, resulting in a time series known as the neural activity index ^115,121^ for each of the 3559 brain sources over five minutes of recording.

To be noted, our analytical choices in source reconstruction further contributed to mitigating volume conduction and source leakage effects. Beamforming implementations, such as the one employed here, are particularly effective for reducing volume conduction by focusing on reconstructing source activity while suppressing signals from unwanted directions. Additionally, we applied regularization to the beamformer weights (see, for example, ^121^), which not only counteracted the bias toward the center of the head but also prevented overfitting and mitigated noise, contributing to address volume conduction issues. Moreover, by prioritizing source-space analyses over sensor-space analyses, we were better able to disentangle distinct sources and limit the effects of volume conduction. We also utilized state-of-the-art algorithms for co-registration of MRI with MEG data. Wherever possible, individual MRIs were employed, and we used the “rhino” algorithm implemented through integration of MATLAB and FSL code for highly effective co-registration. These measures reduced errors in the lead field matrix, further minimizing volume conduction effects and enhancing the accuracy of the analysis.

This whole procedure was applied to PL and RS MEG sensor data, separately.

### Generalized Eigendecomposition

Generalized eigendecomposition (GED) ^30–32,92^ was applied to the 3559 reconstructed brain sources, in order to separate brain networks operating at specific frequencies. Given a frequency of interest, this technique identifies the weighted combination of voxels which best separates narrow oscillatory activity from broadband background activity, by maximizing the contrast between patterns of covariance in input data ^33–35,94^. A major novelty of our approach consists of applying the technique to a highly dense set of reconstructed brain sources, which we argued would yield physiologically interpretable estimations of connectivity within brain networks identified based on a criterion of frequency-specificity.

On a first iteration, we targeted the stimulation frequency of 2.4 Hz, with the goal of identifying brain networks attuned to process the auditory stimulus. The first step consisted of narrow filtering the multivariate signal in voxel space, for which we designed a wavelet kernel as a Gaussian function in the frequency domain, with center at 2.4 Hz and width of 0.3 Hz at half of the maximum gain. The settings were based on previous applications of GED to EEG data in the context of neural entrainment to isochronous auditory stimuli presented at a similar frequency ^33,35^. We then filtered the broadband voxel data via element-wise multiplication between the broadband spectrum and the wavelet kernel in frequency domain, and transformed the resulting tapered spectrum in the time domain via inverse fast Fourier transform.

Subsequently, we repeated the analysis for a large sample of 86 frequencies ranging from 0.2 to 97.6 Hz, equally spaced in intervals of 1.2 Hz for the 80 frequencies above the stimulation frequency and 0.2 for the six frequencies below. In order to compare in the sample elements with and without a harmonic ratio with respect to the stimulation, frequencies were defined as an evenly spaced series where every second number was a multiple integer of the stimulation frequency (harmonic), and an evenly spaced halving series where every second number was a divisor of the stimulation frequency (sub-harmonic). The width of the filters used to compute the narrow signals was optimized for every frequency, on a log10-spaced scale.

For each frequency, the components of interest were separated by applying a spatial filter to the voxels data matrix. In practice, spatial filtering consists of weighting the contribution of all 3559 voxels to the narrow activity. The set of vectors *W* (weights) was calculated by solving the following equation:

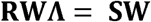

where *R* is the reference covariance matrix of the broadband signal in voxel space and *S* is the covariance matrix of the narrow signal in voxel space filtered at the frequency of interest. GED identifies the set of eigenvectors *W* associated with the eigenvalues Λ, containing the weights to best separate the signal (*S*) covariance from the reference (*R*) covariance matrix. The *S* and *R* covariance matrices were computed in one step via matrix multiplication of the mean-centered narrow and broadband voxel data, respectively.

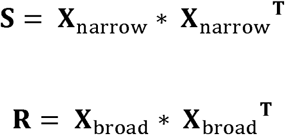

To improve the numerical stability of the GED performance in case of rank deficiency, regularization was applied to both covariance matrices by adding a small constant to their diagonal ^11,122^. For *S*, the constant was 10^-6^, whereas for *R* it was 1% of its average eigenvalues.

The eigenvectors associated to the 10 largest eigenvalues were taken as spatial filters, individually transposed and multiplied by the broadband voxel data matrix to reconstruct the time series of our target signal components. When applying this spatial filter to the voxel data matrix, the resulting signal component is physiologically interpretable as a *network activation time series* (*Y*).

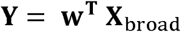

Following the computation of the time series, we computed the spatial projection of the components in voxel space. To do so, we multiplied the raw weights *w* by the *S* covariance matrix and computed its absolute value, obtaining the absolute voxel activation strengths associated with the estimated latent factors (i.e., the networks) ^51^. In order to facilitate comparisons across components and frequencies, we normalized the scale of all activation patterns between 0 and 1 by dividing each vector by its maximum value. The resulting topographical distribution of the components were interpreted as *network activation patterns* (*A*).

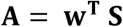

### Randomizations

The entire GED procedure was applied separately to PL and RS reconstructed brain voxel data. In some respects, the resting state worked as physiological control condition to compare the networks’ engagement in auditory processing with the networks in the same frequency band at rest. However, the intrinsic dynamics of the brain at rest are, in principle, still identifiable as frequency-specific networks via GED.

In order to systematically test the sensitivity of FREQ-NESS to the temporal and spatial dimensions of latent brain networks, we independently disrupted their temporal and spatial organization by performing two types of randomizations on our voxel data in the RS condition. The randomization strategies were designed as follows: (1) *label randomization*: the voxel indexes were shuffled row-wise; (2) *point-wise label randomization*: the voxel indexes were shuffled row-wise, independently for each timepoint. Importantly, note that the randomizations were performed after filtering in the narrowband data in order not to alter the filtering procedure. The selective effects of each randomization strategy on the GED performance are presented in **Figure 4**.

Moreover, to statistically assess changes in the RS network landscape resulting from the randomization strategies, we performed a series of Wilcoxon signed-rank tests. Specifically, we compared the original RS data (see **Figure 4a**) with Randomization 1 (Label, see **Figure 4b**) and Randomization 2 (Point-wise label, see **Figure 4c**) across the entire frequency spectrum for the top three eigenvalues. FDR correction was applied to account for multiple comparisons.

### Cross-frequency coupling (CFC)

By their nature, the frequency-resolved network activations estimated via GED are maximally expressed in a defined narrow frequency band. This feature makes cross-frequency coupling (CFC) an optimal approach to investigate interactions across networks since narrowband activity accounts for the largest part of the variance in each network. For each frequency, the first component was narrowband filtered using the same parameters previously defined for GED. Subsequently, the narrowband component was Hilbert-transformed to produce an analytic signal.

The first two components attuned to the stimulation frequency at 2.4 Hz were processed as low-frequency modulators. The phase time series were extracted from the analytic signal and divided into 36 bins (bin size = 10°) ^54^. For each frequency, the first component was processed as a carrier signal. Power time series were computed as the squared magnitude of the analytic signal, and the mean power within each phase bin was computed to produce power distributions over the modulators’ phase ^56^. Finally, a sine function was fitted to each distribution using the sineFit MATLAB function ^123^, and its amplitude was used as an index of modulation strength ^34^.

### Statistical testing

Signed-rank tests were performed to test for significant differences in the network landscapes across experimental conditions. A two-tailed test of the eigenvalues was performed testing PL against RS, for each component and each frequency. For this test, we assumed that significant changes could be found in both directions, due to the possible attunement or suppression of frequency-specific brain networks resulting from the auditory stimulation.

A right-tailed test of the strength of the modulation by the low-frequency auditory components was performed testing PL against RS. The test was repeated for the first and the second low-frequency modulators, for each component and each frequency. For this test, we hypothesized that PL would result in stronger modulation because no auditory network was expected to be attuned to the stimulation frequency in Rest.

False-discovery rate (FDR) correction was applied to all statistical tests to define a significance threshold adjusted for multiple comparisons. Furthermore, to complement the original FDR-corrected statistical tests, we conducted an additional analysis using a cluster- based permutation test with Monte-Carlo simulations (MCS) to correct for multiple comparisons.

## Data availability

The pre-processed neuroimaging data generated in this study will be made available upon reasonable request.

## Code availability

The MEG data were first pre-processed using MaxFilter 2.2.15. Then, the data were further pre-processed using Matlab R2016a or later (MathWorks, Natick, Massachusetts, United States of America). Specifically, we used codes from the Oxford Centre for Human Brain Activity Software Library (OSL), FMRIB Software Library (FSL) 6.0, SPM12, and Fieldtrip. The full analysis pipeline used in this study as well as the code developed for FREQ-NESS are available in the following GitHub: https://github.com/mattiaRosso92/Frequency-resolved_brain_network_estimation_via_source_separation_FREQ-NESS

## Acknowledgements

The Center for Music in the Brain (MIB) is funded by the Danish National Research Foundation (project number DNRF117).

M.R. is supported by IPEM Institute for Systematic Musicology, Ghent University, Belgium, with a Methusalem grant awarded by the Flemish Government, Belgium, to Professor Marc Leman.

L.B. is supported by Lundbeck Foundation (Talent Prize 2022), Carlsberg Foundation (CF20-0239), Center for Music in the Brain, Linacre College of the University of Oxford and Nordic Mensa Fund.

M.L.K. is supported by Center for Music in the Brain and Centre for Eudaimonia and Human Flourishing, which is funded by the Pettit and Carlsberg Foundations.

We thank Simjon Radloff, Anastasiia Popova and Antonieta Martínez-Guerrero for their extensive and tireless work during the data collection.

## Author contributions statement

M.R. and L.B. conceived the hypotheses. L.B. and E.B. designed the experiment and collected the data. L.B., M.L.K., E.B., and P.V. recruited the resources for the experiment. M.R. and L.B. performed pre-processing, signal processing, statistical analysis and implemented the code for the FREQ-NESS analysis pipeline. P.K., M.L.K., E.B., and P.V. provided essential help to interpret and frame the results within the neuroscientific literature. M.R., G.F.R. and L.B. wrote the first draft of the manuscript. G.F.R., M.R., and L.B. prepared the figures. All the authors contributed to and approved the final version of the manuscript.

## Competing interests statement

The authors declare no competing interests.

## Notes

### Competing Interest Statement

The authors have declared no competing interest.

### Summary of Updates

We have revised the text to enhance clarity, readability, and alignment with previous literature. Additionally, we have conducted further analyses that largely confirm our original results while providing additional technical details and insights into our robust methodology.

